# Integrative analysis of genomic variants reveals new associations of candidate haploinsufficient genes with congenital heart disease

**DOI:** 10.1101/2020.06.25.169573

**Authors:** E Audain, A Wilsdon, J Breckpot, JMG Izarzugaza, TW Fitzgerald, AK Kahlert, A Sifrim, F Wünnemann, Y Perez-Riverol, H Abdul-Khaliq, M Bak, AS Bassett, JW Belmont, DW Benson, F Berger, I Daehnert, K Devriendt, S Dittrich, P Daubeney, V Garg, K Hackmann, K Hoff, P Hofmann, G Dombrowsky, T Pickardt, U Bauer, BD Keavney, S Klaassen, HH Kramer, CR Marshall, DM Milewicz, SA Lemaire, J Coselli, ME Mitchell, A Tomita-Mitchell, SK Prakash, K Stamm, AFR Stewart, CK Silversides, R Siebert, B Stiller, JA Rosenfeld, I Vater, AV Postma, A Caliebe, JD Brook, G Andelfinger, ME Hurles, B Thienpont, LA Larsen, MP Hitz

## Abstract

Congenital Heart Disease (CHD) affects approximately 7-9 children per 1000 live births. Numerous genetic studies have established a role for rare genomic variants at the copy number variation (CNV) and single nucleotide variant level. In particular, the role of *de novo* mutations (DNM) has been highlighted in syndromic and non-syndromic CHD. To identify novel haploinsufficient CHD disease genes we performed an integrative analysis of CNVs and DNMs identified in probands with CHD including cases with sporadic thoracic aortic aneurysm (TAA). We assembled CNV data from 7,958 cases and 14,082 controls and performed a gene-wise analysis of the burden of rare genomic deletions in cases versus controls. In addition, we performed mutation rate testing for DNMs identified in 2,489 parent-offspring trios. Our combined analysis revealed 21 genes which were significantly affected by rare genomic deletions and/or constrained non-synonymous *de novo* mutations in probands. Fourteen of these genes have previously been associated with CHD while the remaining genes (*FEZ1, MYO16, ARID1B, NALCN, WAC, KDM5B* and *WHSC1*) have only been associated in singletons and small cases series, or show new associations with CHD. In addition, a systems level analysis revealed shared contribution of CNV deletions and DNMs in CHD probands, affecting protein-protein interaction networks involved in Notch signaling pathway, heart morphogenesis, DNA repair and cilia/centrosome function. Taken together, this approach highlights the importance of re-analyzing existing datasets to strengthen disease association and identify novel disease genes.

## Introduction

Congenital Heart Disease (CHD) accounts for a large fraction of foetal and infant deaths, with incidence rates ranging from 7-9 per 1000 live births^1^. Within the last 30 years, survival rates have substantially increased due to improvements in surgical, interventional and clinical intensive care resulting in a rapidly growing number of CHD survivors reaching adulthood^2^. Nevertheless, there is still increased morbidity and mortality in individuals with CHD, resource utilization is high especially among severely affected patients, and importantly, the underlying etiology remains unclear for the majority of cases.

CHD is multifactorial, with both environmental and genetic risk factors^3,4^. Familial aggregation of CHD including Thoracic aortic aneurysm (TAA), as well as a large proportion of genomic copy number variants (CNVs) and *de novo* intragenic mutations (DNMs) in probands with CHD suggest a strong genetic component. An estimated 4-20% of CHD cases are due to rare CNVs, suggesting that a significant part of CHD is caused by gene-dosage defects^5^. Recently, exome sequencing in large cohorts has been used to identify novel disease genes and strengthen known disease associations through the demonstration of an excess of *de novo* protein truncating variants (PTV) and rare inherited loss-of-function (LOF) variants in probands with CHD^6,7^.

Overlaying both CNVs and PTVs has been used to define novel CHD relevant disease genes in contiguous gene disorders^8,9^. Following this principle, we have performed a genome-wide integrative meta-analysis of published and publicly available datasets of CNVs and DNMs identified in probands with CHD. This analysis, which is one of the larger meta-analyses of genomic variants in CHD so far, strengthens the disease association of known CHD genes and identifies novel haploinsufficient CHD candidate genes.

## Results

### Cohort description and workflow

We assembled a cohort with 7,958 cases (comprising both non-syndromic CHD, syndromic CHD and TAA cases) and 14,082 controls (**Supplemental Table 1**). Of the total of cases, 777 (∼10%) were diagnosed with Thoracic Aortic Aneurysm (TAA). An overview of the sources used to assemble the present cohort is listed in **Supplemental Table 2** (for CHD cases) and **Supplemental Table 3** (for controls). We applied a set of quality control filters to our assembled CNV data before performing case-control association tests (see Methods section). In addition, common CNVs (minor allele frequency (MAF) in controls > 0.01) were excluded from the analysis. After filtering, 6,746 cases and 14,024 controls remained for further downstream analysis. Furthermore, we built a dataset of *de novo* mutations (DNMs) identified in 2,489 probands with CHD from parent-offspring trios^6,7^.

### CNV burden test of known CHD genes

Haploinsufficiency has been shown to cause a reasonable proportion of CHD^5^. Thus, genes known to be associated with CHD and genes which are intolerant for LOF mutations should be deleted more often in probands with CHD than in controls. To test this hypothesis, we performed a CNV burden test using sets of genes known to be involved in CHD. In addition, we included genes known to be associated with developmental disorders, a curated list of known haploinsufficient disease genes^10^, autosomal recessive disease genes^11,12^ and genes predicted to be intolerant to LOF mutations (based on the observed/expected LOF ratio from gnomAD^13^). The burden test was performed using a logistic regression framework^14^ (implemented in PLINK v1.7), where the phenotype is regressed on the number of genes deleted and covariates (see Methods). **Figure 1A** (extended **Supplemental Table 4**) summarizes the results from the burden test on the different gene sets: known CHD genes (grouped in syndromic, non-syndromic, monoallelic and biallelic), developmental disorder genes, haploinsufficiency disease genes, autosomal recessive genes and all protein coding genes. We tested all protein coding genes to address the possibility that the analyses could be biased by differences in the CNV rate within the case and control groups, since we have assembled our cohort from different datasets. We did not observe genome-wide (all tested protein coding genes) enrichment (*P* = 0.39, *OR* = 0.99) nor enrichment in the autosomal recessive gene set (*P* = 0.52, *OR* = 1.03) when comparing rare CNV deletions in cases vs controls. In contrast, the analysis revealed significant differences in the burden of CNV deletions between cases and controls for the set of haploinsufficiency genes (*P* = 8.29 × 10^−13^, *OR* = 2.27). As expected, our analysis revealed significant enrichment for the set of known CHD genes, which is mainly explained by the contribution of monoallelic CHD genes (*P* = 2.04 × 10^−31^, *OR* = 4.13) and syndromic CHD gene set (*P* = 1.66 × 10^−33^, *OR* = 4.06). Unlike the monoallelic and syndromic CHD gene sets, no significant enrichment was found for the nonsyndromic (*P* = 0.75, *OR* = 1.16) and biallelic (*P* = 0.08, *OR* = 1.87) CHD gene sets. Our analysis revealed a moderate enrichment of rare CNVs in the developmental disorder gene set (*P* = 6.90 × 10^−11^, *OR* = 1.75).

**Figure 1.**
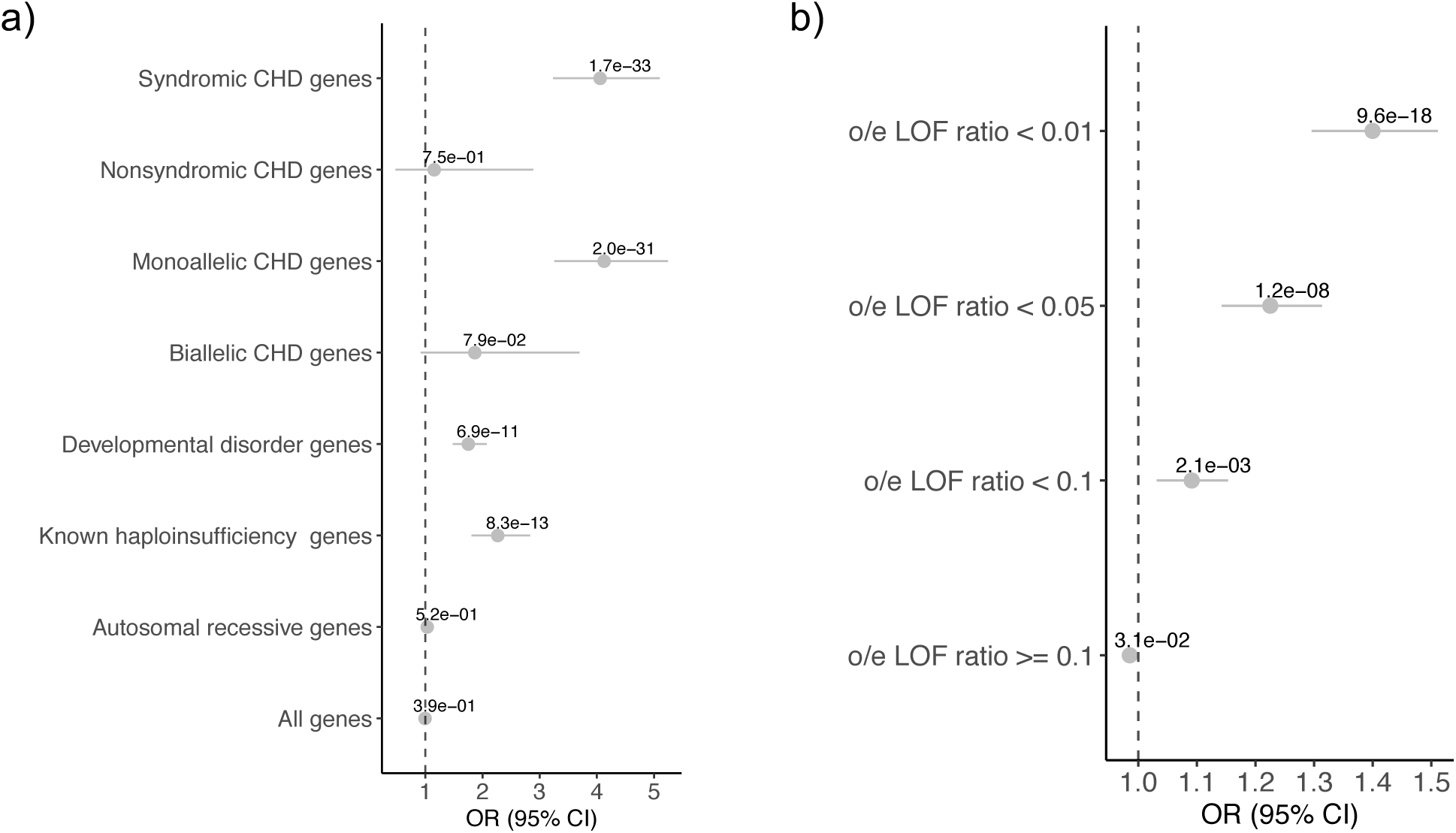
CNV burden test on known gene sets (a) and constraint LOF genes at different observed/expected LOF ratio thresholds (b). The forest plot shows the odds ratio (dots), the 95% confidence intervals indicating the certainty about the OR (interrupted line) and the *P-value* in the indicated gene sets.

When the regression-based analysis was performed at different levels of the observed/expected LOF ratio (*oeLOF*) constraint metric (**Figure 1B, Supplemental Table 5**), we observed the higher enrichment toward the most LOF constrained genes (*oeLOF* < 0.01, *P* = 9.55 × 10^−18^, *OR* = 1.40) and still a moderate enrichment for genes with *oeLOF* < 0.1 (*P* = 0.002, *OR* = 1.09). No enrichment was observed in the set with *oeLOF* ratio >= 0.1 (*P* = 0.03, *OR* = 0.99). Based on these results we conclude that haploinsufficiency causes a significant component of CHD.

### Genome-wide identification of haploinsufficiency candidate disease genes for CHD

To perform a systematic, genome-wide identification of potential haploinsufficient CHD disease genes and loci, we analysed the CNV burden of 19,969 protein coding genes (GENCODE v19). To this end, we compared the number of rare CNV deletions (MAF < 0.01) among cases and controls for each gene and identified genes with significant CNV burden using a permutation test (significance level of adjusted *P* < 0.05, see Methods). The distributions of rare CNV deletions in CHD cases across all 22 human autosomes is shown in **Figure 2**. Significant candidate genes had a median number of 12 overlapping CNVs in cases, compared to a median of 0 overlapping CNVs in controls (**Supplemental Figure 1a**). Because CNVs can be large chromosomal aberrations, multiple genes were affected by some of the CNVs. In total, 528 genes (**Supplemental Table 6**) reached significance (Permutation test, *P* adjusted < 0.05). These 528 genes encompass a total of 63 loci (**Supplemental Table 6**, highlighted in magenta in **Figure 2**). The sizes of these loci range from 558 bp to 10.5 Mbp, with a median value of 243 Kbp (**Supplemental Figure 1b**). The number of genes per locus ranged from 1 to 48, with a median value of 3 (**Supplemental Figure 1c**). Only 16 loci contained a single gene (**Supplemental Table 6**). In addition, we tested previously described CNV deletion syndrome regions (https://decipher.sanger.ac.uk/disorders#syndromes/overview) associated with developmental disorders and/or CHD for enrichment in our analysis (**Methods**). We found eight of these regions enriched in the dataset (**Supplemental Table 7**), with the 16p11.2-p12.2 locus being the region with the largest number of deletions in cases (n=230).

**Figure 2.**
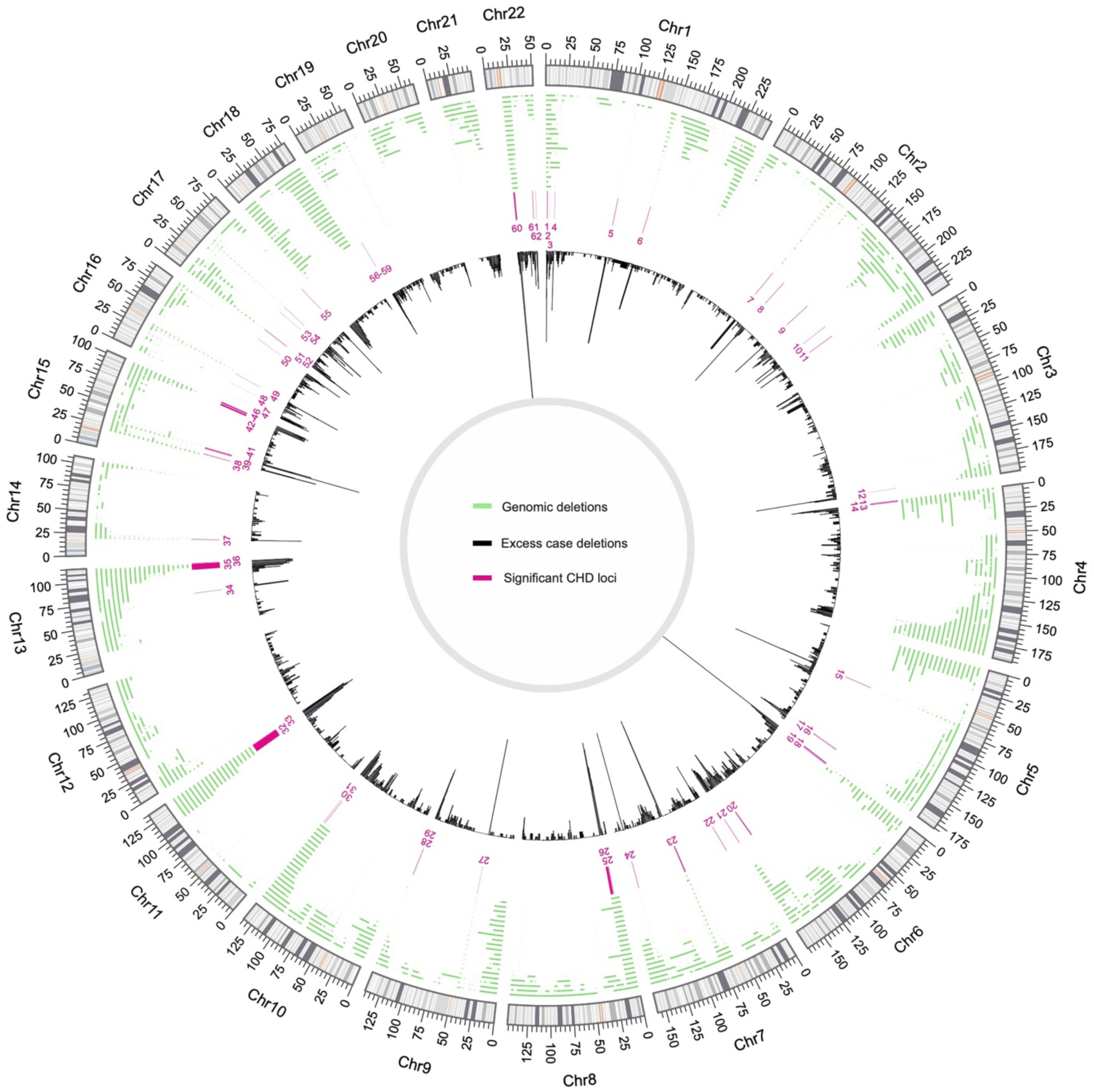
CNV deletion distribution across the 22 autosomes. The plot shows the distribution of rare CNV deletions (green track) in CHD cases, the differences between the overlapping CNV deletions in cases and controls (black track) and highlight the location of the 63 significant loci discovered (in magenta).

### Shared genetic architecture of CHD and TAA

We independently performed a genome-wide test without the TAA cases to evaluate its impact on CHD. As expected, most of the genes (447 out of 528) remained significant after removing the contribution of TAA cases, since ∼90% of the cases in the analyzed CNV cohort were CHD. Ten genes were significantly enriched independently when analyzing CHD and TAA cases, while 61 were significantly enriched only in TAA cases (**Supplemental Table 8**).

### De novo *mutation analysis*

To identify an independent set of haploinsufficient CHD candidate genes, we combined *de novo* mutations identified in two large-scale CHD case-control studies^6,7^ and performed a gene-based *de novo* mutation (DNM) burden test^15^. We analysed a total of 4,192 rare DNMs within 2,534 genes in the patient cohort. After classifying every variant into functional groups (see Methods section), 526 of these variants were predicted to be protein-truncating and, 2,647 were missense. We evaluated for potential differences of the DNM rates between cohorts (see Methods). Comparison of the rate of each variant type across the groups was non-significant (*P* > 0.05, Poisson test, **Supplemental Table 9**).

We used two available statistical methods, Mupit^15^ and DeNovoWEST^16^, which test the significance of observed DNM at gene level, by comparing the number of observed mutations with the number of expected mutations (based on a sequence-dependent mutation recurrence rate, see Methods). While Mupit focuses on enrichment of LOF DNMs specifically, the DeNovoWEST test incorporates missense constraint information at variant level and applies a unified severity scale at variant level based on the empirically-estimated positive predictive value of being pathogenic. Based on the complementary results of both tests^16^, we reported the minimal observed DNM *p-value* (*P*_*dnm*_) per gene.

We identified 14 genes significantly enriched in the DNM analysis (*P* < 0.05 after Bonferroni correction for multiple testing, **Supplemental Table 10**). All of these genes were affected by at least two constrained non-synonymous DNM (nsDNM) and show significant overlap with 11/14 (78.6%) of the genes being known CHD disease genes. *CHD7* (OMIM 214800) was the most significant haploinsufficient gene (*P* = 2.84 × 10^−26^) with 18 nsDNM identified in the patient cohort. Other highly enriched genes for nsDNM - *KMT2D* (OMIM 147920)^17^, *KMT2A* (OMIM 605130)^18^, *NSD1* (OMIM 117550)^19^, *TAB2* (OMIM 614980) ^20^, and *ADNP* (OMIM 615873) ^21^ - have been previously associated with different types of neurodevelopmental disorders with co-occurrence of CHD. In the case of *KDM5B* (OMIM 618109), it has only been described in the context of a recessive neurodevelopmental phenotype with cases presenting ASD (Atrial septal defects) ^22,23^.

We next evaluated the distribution of o/e LOF ratio at different levels of DNM enrichment (genes were split based on *P*_*dnm*_). Since the o/e ratio of LOF variation in each gene is strongly affected by its length, we instead used the 90% upper bound of its confidence interval (termed LOEUF), which keeps the direct estimate of the o/e ratio and allows to distinguish small genes from large genes, as suggested by Karczewski et al^13^. We observed that the genes with higher enrichment for nsDNM (lower *P*_*dnm*_) show a significant decreased LOEUF compared to the mean of all protein coding genes (**Figure 3**).

**Figure 3.**
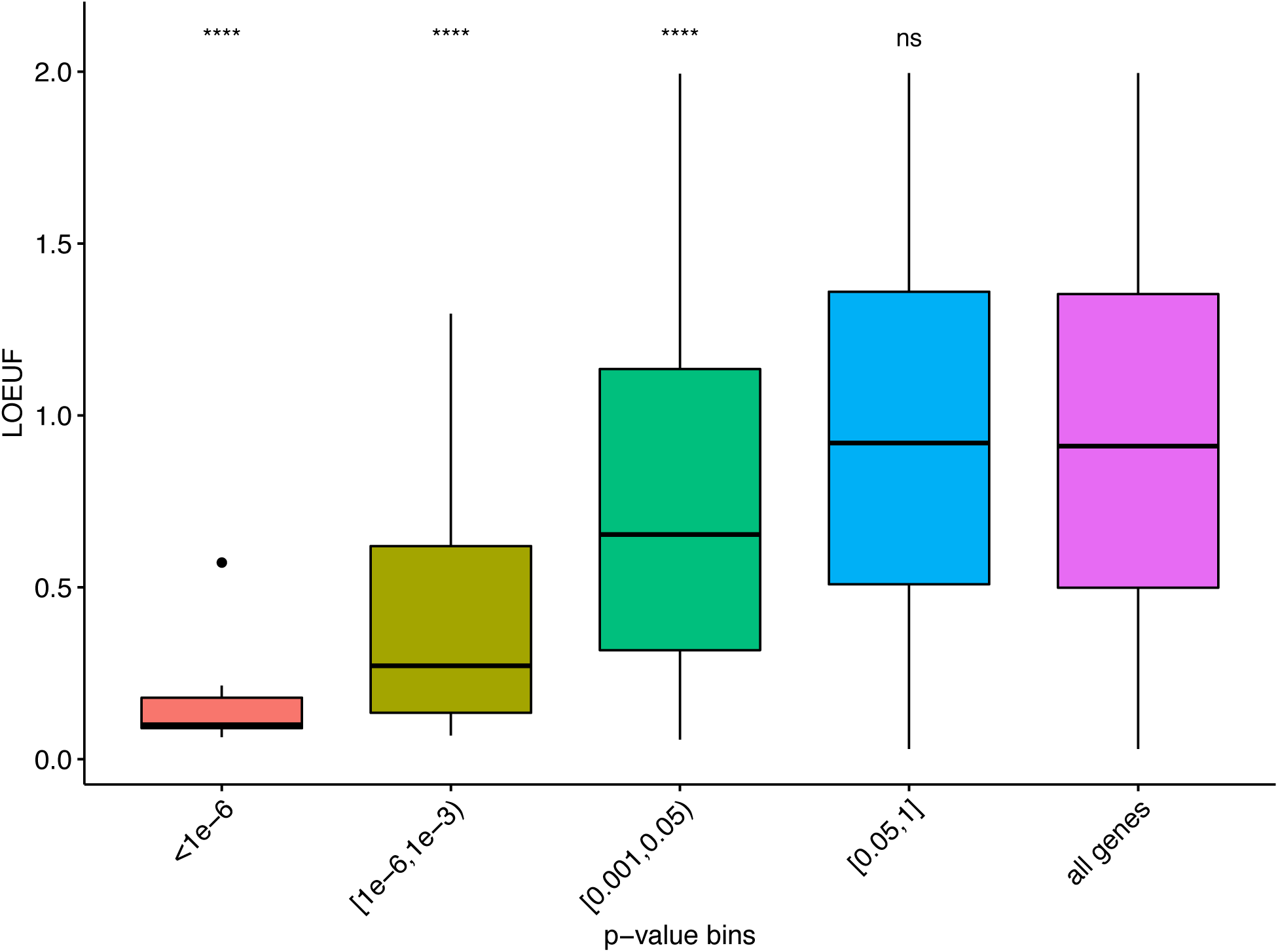
Comparison of the distribution of LOEUF metric at different level of significance of nsDNM-enriched genes. X-axis denotes the P-values from the DNM analysis (binned). Y-axis denotes the o/e LOF ratio upper bound fraction (LOEUF). All groups were compared against the LOEUF distribution of all protein coding genes (purple). Differences between the distributions were tested using a two-sided Wilcoxon rank sum test. ****: P<0.0001, ns: non-significant.

### Integration of DNM and CNV results

To identify high confidence haploinsufficient CHD disease genes, we performed a joint analysis integrating the results from the CNV and the DNM analysis. We combined the results from both analyses (*P*_*dnm*_ and *P*_*cnv*_) using the Fisher combine method. We demonstrated that both enriched genes for DNM and CNV deletions are significantly represented among LOF constraint genes (measured by the o/e LOF ratio). Therefore, we applied a Bonferroni multiple testing correction using independent hypothesis weighting^24^ (IHW) by incorporating the gene o/e LOF ratio, as a measure of haploinsufficiency (**Supplemental Figure 2**). Our analysis revealed 21 genes that were significantly enriched for CNV deletions and/or non-synonymous DNM (**Table 1**). A gene was included in the final set of haploinsufficient CHD disease genes if it reached a significant corrected *metaP* < 0.05 (after Bonferroni adjustment with IHW).

**Table 1.** Top 21 significant genes arising from both the permutation-based test and the DNM rate-based test. Cases/Controls: Number of cases and controls carrying CNV deletions overlapping the gene in the CNV analysis. *P*_*cnv*_: p-value from the CNV permutation test. nsDNM: Number of constrained non-synonymous mutations in the *de novo* analysis. *P*_*dnm*_: p-value from the DNM analysis. Significant: The analysis where the gene was significant (dnm: DNM analysis, cnv: CNV analysis, both: Both analysis, none: Non-significant neither DNM nor CNV analysis). *metaP*: combined p-value (*P*_*dnm*_ and *P*_*cnv*_) using the Fisher method. *P*_*ihw*_: Bonferroni corrected p-value using independent hypothesis weighting (IHW) and LOEUF metric as covariate. LOEUF: o/e LOF ratio upper bound fraction from gnomAD. *All the 21 genes were significant after combining their p-values and applying Bonferroni correction. ^1^Evidence is from mouse models^35,36^.

| Gene | CNV |  |  | DNM |  | *Significant | Combined |  | LOEUF | Known CHD |
| --- | --- | --- | --- | --- | --- | --- | --- | --- | --- | --- |
| | case | control | $P_{cnv}$ | nsDNM | $P_{dnm}$ | | $metaP$ | $P_{ihw}$ | | |
| **CHD7** | 6 | 1 | 6.80E-03 | 18 | 2.84E-26 | dnm | 1.25E-26 | 8.05E-23 | 0.076 | Yes |
| **KMT2D** | 0 | 0 | 1.00E+00 | 18 | 1.32E-25 | dnm | 7.67E-24 | 4.93E-20 | 0.103 | Yes |
| **NSD1** | 1 | 1 | 5.63E-01 | 12 | 1.00E-14 | dnm | 1.90E-13 | 2.14E-09 | 0.095 | Yes |
| **KMT2A** | 0 | 0 | 1.00E+00 | 7 | 1.00E-14 | dnm | 3.32E-13 | 1.86E-09 | 0.065 | Yes |
| **NOTCH1** | 10 | 24 | 1.00E+00 | 7 | 1.00E-14 | dnm | 3.32E-13 | 2.14E-09 | 0.097 | Yes |
| **TAB2** | 12 | 0 | 1.00E-04 | 5 | 3.46E-09 | both | 1.03E-11 | 5.75E-08 | 0.098 | Yes |
| **ANKRD11** | 13 | 0 | 1.00E-04 | 3 | 2.32E-05 | cnv | 4.85E-08 | 2.72E-04 | 0.107 | Yes |
| **WHSC1** | 11 | 0 | 1.00E-04 | 3 | 8.73E-05 | cnv | 1.71E-07 | 9.96E-04 | 0.119 | No<sup>1</sup> |
| **ADNP** | 0 | 0 | 1.00E+00 | 4 | 9.94E-09 | dnm | 1.93E-07 | 1.13E-03 | 0.123 | Yes |
| **DYRK1A** | 4 | 0 | 1.43E-02 | 4 | 9.46E-07 | dnm | 2.59E-07 | 1.64E-03 | 0.214 | Yes |
| **NALCN** | 10 | 1 | 1.00E-04 | 3 | 1.76E-04 | cnv | 3.32E-07 | 6.83E-03 | 0.522 | No |
| **ELN** | 30 | 0 | 1.00E-04 | 2 | 1.77E-04 | cnv | 3.34E-07 | 7.50E-03 | 0.871 | Yes |
| **WAC** | 7 | 0 | 4.00E-04 | 3 | 1.31E-04 | none | 9.33E-07 | 5.44E-03 | 0.084 | No |
| **RBFOX2** | 1 | 0 | 3.45E-01 | 4 | 1.59E-07 | dnm | 9.72E-07 | 6.25E-03 | 0.194 | Yes |
| **KANSL1** | 94 | 110 | 2.00E-04 | 2 | 3.38E-04 | none | 1.19E-06 | 6.92E-03 | 0.238 | Yes |
| **MYO16** | 13 | 2 | 1.00E-04 | 2 | 8.38E-04 | cnv | 1.45E-06 | 1.74E-02 | 0.272 | No |
| **MED13L** | 2 | 1 | 2.70E-01 | 4 | 4.58E-07 | dnm | 2.09E-06 | 1.22E-02 | 0.064 | Yes |
| **KDM5B** | 0 | 0 | 1.00E+00 | 4 | 1.45E-07 | dnm | 2.43E-06 | 4.97E-02 | 0.572 | No |
| **GATA6** | 0 | 0 | 1.00E+00 | 5 | 1.80E-07 | dnm | 2.98E-06 | 1.92E-02 | 0.174 | Yes |
| **ARID1B** | 4 | 0 | 1.31E-02 | 4 | 1.48E-05 | none | 3.20E-06 | 1.87E-02 | 0.102 | No |
| **FEZ1** | 10 | 0 | 1.00E-04 | 2 | 2.22E-03 | cnv | 3.62E-06 | 4.30E-02 | 0.414 | No |

### Significant CHD genes are highly and/or differentially expressed in the heart

We next evaluated the expression pattern of the 21 significant genes (Bonferroni corrected *metaP* < 0.05) in the heart using RNA-Seq data from human tissues at different developmental time points^25^. We stratified the analysis based on stages of heart development (see Methods). Our analysis revealed that the most significant genes (*metaP* < 1×10e^-5^) show significantly increased mean expression in the heart (*P* < 0.0001, Wilcox test) at different developmental stages (development, maturation and infant/adult), compared to all protein coding genes (**Figure 4**). Moreover, 18 out of 21 genes fall in the in the top quartile of heart expression in both developmental and maturation stages, while 8/21 genes remain highly expressed in adulthood (**Supplemental Figure 3**). To complement our expression analysis, we compared gene expression during human heart development with expression in two other mesodermal organs: kidney and liver. This allowed identification of genes with significant changes in its expression levels during crucial heart developmental stages, which would have not been possible when focusing on expression levels alone (**Methods**). We found that 17 out of 21 CHD candidate genes are differentially expressed in the heart (*R*^*2*^ > 0.50, Bonferroni corrected *P* < 0.01) when compared to its expression levels in kidney and/or liver. Interestingly, the three genes (*FEZ1, NALCN* and *MYO16*) which are not among the highly expressed genes, were found to be significantly differentially expressed during heart development compared to kidney and/or liver (**Supplemental Table 11**).

**Figure 4.**
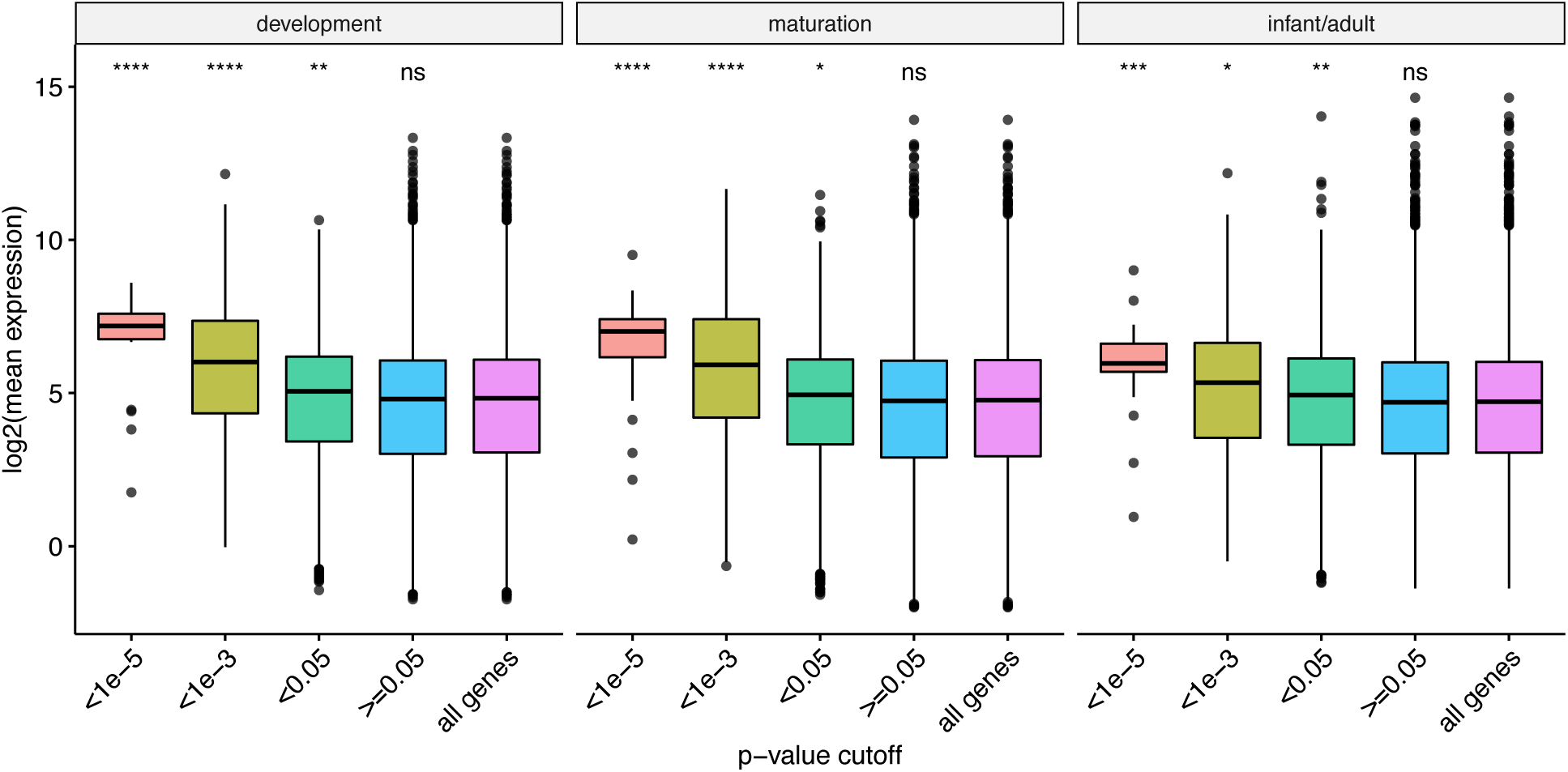
Comparison of the mean expression (heart) distribution at different *metaP* cut-offs. Panels show three different heart development stages: early development, maturation and infant/adult. X-axis denotes the combined p-value from DNM and CNV analysis (*metaP*, at different cut-offs). Y-axis denotes the genes’ mean expression in the heart (log scale). The 21 significant candidate CHD genes (Table 1) are contained in the fraction with the higher expression (red box). Differences between the distributions were tested using a two-sided Wilcoxon rank sum test (reference group: all genes). ****: P<0.0001, ***: P<0.001, **: P<0.01, *: P<0.05, ns: non-significant.

### CNV/DNMs hinder the function of specific protein complexes

CNVs and DNMs can affect heart development either through haploinsufficiency of a single gene, or through its combined impact on the function of several genes. Indeed, oligogenic models have been implicated in CHD, and proteins acting in the same complex or pathway are known to be encoded in genomic clusters^26,27^. We therefore conducted a systems-level analysis to identify global mechanisms by which haploinsufficiency might promote CHD. In particular, we assessed the combined effect of CNVs and DNMs with respect to human protein-protein interactions (PPIs). The InWeb^28^ and ConsesusPathDB^29^ databases provides ranked information about experimentally determined physical interactions and, therefore, serves as a proxy to understand the functional effects of CNV/DNMs on human protein complexes. The genes with Benjamini–Hochberg adjusted *metaP* < 0.05 (n=492 genes) were used as seeds to build a PPI network from the data available in InWEb and ConsessusPathDB. No additional interections were considered. The final network consisted of 164 proteins and 290 interactions (**Supplemental Figure 4**). A total of 10 overlapping sub-clusters within this network were identified using the in-built clustering algorithm implemented in GeNets^30^ (Methods). Gene-ontology (GO) enrichment analysis suggested that four out of these ten sub-clusters are enriched for genes involved in Notch signaling pathway, cardiocyte differentiation, DNA repair and centrosome function (**Figure 5**). All the four clusters accommodate more CNV deletions in CHD cases compared to controls. Six out of the ten sub-clusters did not show significant enrichment for any particular biological process.

**Figure 5.**
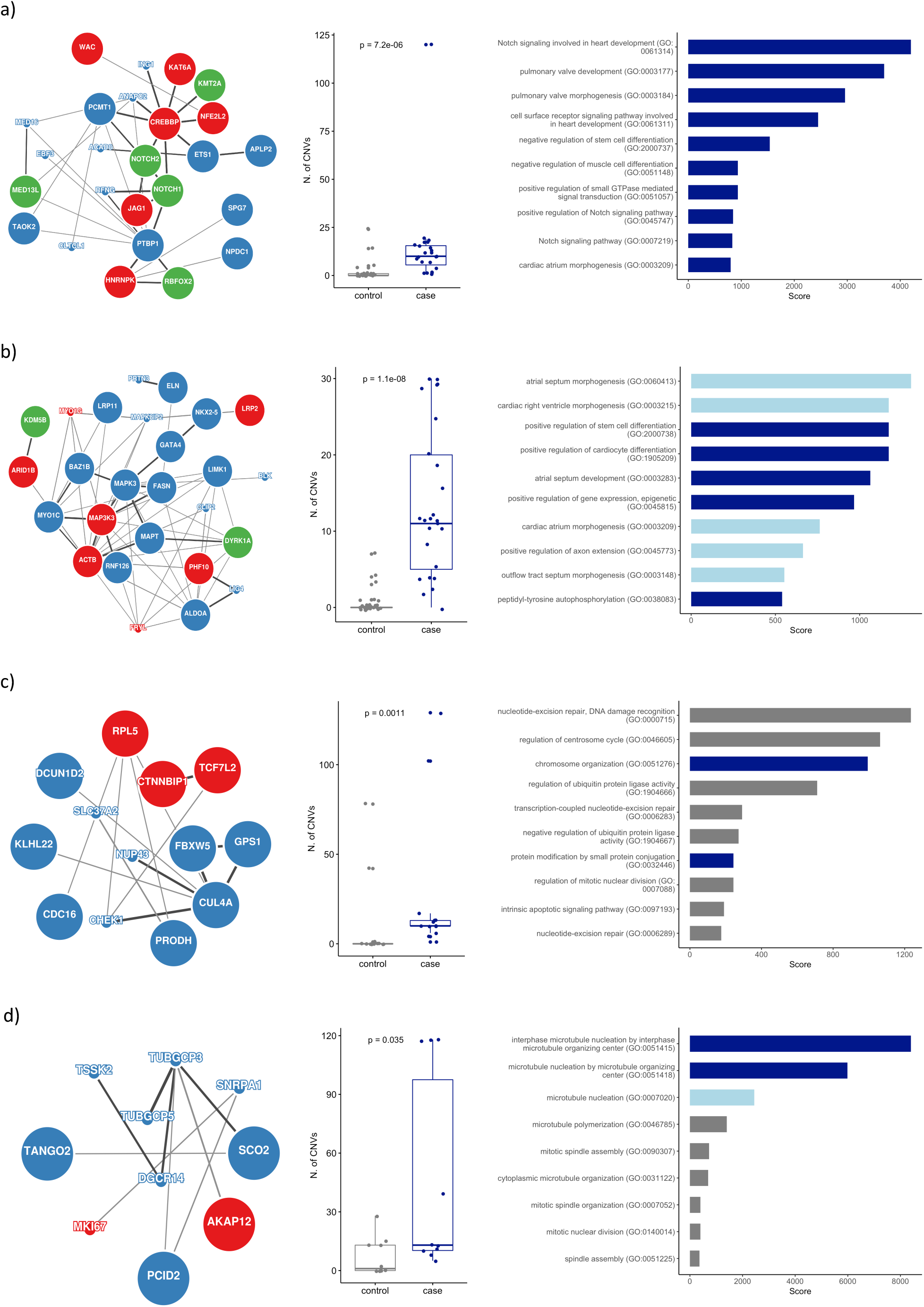
Identification of functional networks enriched for proteins encoded by genes affected by CNVs and/or DNMs associated with CHD. The protein-protein interaction networks (a-d, for clusters 1, 3, 8 and 9 respectively) were identified using GeNets (extended **Supplemental Figure 4**). Proteins are shown as nodes, interactions as edges. Enrichment for CNVs (blue) and DNMs (green) are highlighted. Proteins with no specific enrichment for CNV and/or DNMs but with B-H adjusted *metaP* < 0.05 are highlighted in red. The size of the circles denotes if the genes was found significantly highly and/or differentially expressed in the heart (large circles: significant expression; small circles: non-significant). The distribution of CHD case-CNVs and control-CNVs are shown for each cluster. Significant difference in the CNV distribution was calculated using a Wilcox rank sum test. The horizontal bar plots show the top ten GO enriched terms for each cluster (output from enrichr tool). Bar color encoded the GO biological process significant level (dark blue: FDR < 5%, light blue: FDR 5-10%, grey: FDR > 10%).

## Discussion

We performed a meta-analysis of rare genomic variants in a cohort of 10,447 CHD probands, which provides a useful resource for interpreting CNVs and DNMs identified in patients with CHD. We implemented a statistical approach which allows the integration of different types of genomic variants to discover novel genes associated with CHD. Our data-driven integrative analysis took into account three major criteria at the genomic level: a) gene enrichment for DNM, b) gene enrichment for CNV deletions and c) gene intolerance for LOF mutations. Our analysis identified 21 significant haploinsufficient CHD genes. Fourteen of these are known CHD genes, and the remaining seven genes have not previously been associated with CHD (**Table 1**).

To further strengthen associations, we made use of a newly published human transcriptome atlas covering different developmental, maturation and adult stages in numerous organs^25^. Similar to previous results^7^, our analysis highlights that the majority of the 21 significant genes are highly expressed during critical stages of heart development. Unlike earlier studies^7,31^ which did not address the importance of expression changes over time, we evaluated the differential expression patterns of genes by comparing levels of expression in the heart, kidney and liver at different time points in development. This analysis allowed us to strengthen disease association for genes not falling under the high expression group and highlight the critical importance of all 21 genes independently of the genomic approach. This aspect is complemented by the fact that the majority of genes (14/21) were already known to cause CHD.

Among the 21 likely haploinsufficient disease genes for which the combined analyses showed enrichment (Bonferroni corrected *metaP* < 0.05), 14 genes (*CHD7, KMT2D, KMT2A, NOTCH1, NSD1, TAB2, ANKRD11, ADNP, DYRK1A, RBFOX2, KANSL1, ELN, MED13L* and *GATA6*) are well-established CHD genes, and our data confirms this association. To the best of our knowledge, association between CHD and seven genes (*KDM5B, WHSC1, WAC, NALCN, ARID1B, FEZ1* and *MYO16)* had either not been established, or had been reported in small cases studies or a single individual only.

*KDM5B* is not an established CHD gene thus far, although one patient with compound heterozygous frameshift variants had an ASD^22^. While some have argued against the haploinsufficiency of the gene^32^, our analysis suggests *KDM5B* as a plausible haploinsufficient CHD gene. Additional functional studies are warranted to confirm its role in CHD.

A recent CNV meta-analysis^33^ based on non-syndromic CHD patients found that duplication of *WHSC1* (also known as *NSD2*) is a possible cause of CHD. However, haploinsufficiency of *WHSC1* has not previously been associated with CHD. In support of its role in CHD, *Whsc1* has been reported to cause heart malformations in mouse models^34–36^. In addition, *WHSC1* is known to interact with *NKX2*.5^36^. In spite of this, the low incidence of CHD in individuals with Wolf-Hirschhorn syndrome suggests that haploinsufficiency of *WHSC1* alone does not cause CHD.

Heterozygous truncating mutations in *WAC*, as well as CNV deletions involving this gene, have been recently associated with the DeSanto-Shinawi syndrome, a rare neurodevelopmental disorder characterized by global developmental delay^37,38^. Furthermore, in two non-consanguineous unrelated individuals with heart malformations, among other disorders^39^, microdeletions at 10p11.23-p12.1 (overlapping *ARMC4, MPP7, BAMBI* and *WAC*) were identified. Despite these isolated reports, no definite association between *WAC* and CHD has been established.

DNMs in *NALCN* have been reported to cause a dominant condition characterized by multiple features including developmental delay, congenital contractures of the limbs and face and hypotonia^40,41^. However, among the phenotypes observed, CHD have been not described thus far.

Heterozygous mutation of *ARID1B* is a frequent cause of intellectual disability^42,43^. A recent analysis of 143 patients with *ARID1B* mutations showed that individuals display a spectrum of clinical characteristics. Congenital heart defects were observed in 19.5% of the patients^44^.

*FEZ1* is a neurodevelopmental gene, which has been associated with schizophrenia^45^. *Fez1* has been reported to be regulated by *Nkx2-5* in heart progenitors in mice, suggesting a possible role in heart development^46^.

*MYO16* (*NYAP3*) encodes an unconventional myosin protein, involved in regulation of neuronal morphogenesis^47^. We have not found an association between *MYO16* and heart development in the literature.

Although, several genes have been shown to be altered in syndromic and non-syndromic cases with CHD and TAA (e.g. *HEY2* (manuscript under review), *MYH11*^48,49^, *NOTCH1*^50^, among the 10 genes significant in our analysis for TAA, CHD and the combined scenario, none has been reported previously to be associated with either CHD or TAA. Given the limited data size and only accessing CNV calls from TAA cases, future studies looking at CNV and DNM in both phenotypes are required to establish stronger genotype-phenotype correlation to better understand a possibly shared genetic architecture for the two disease entities.

In addition to the gene-centered analysis, we also applied a systems-level analysis in order to identify potential novel pathophysiological mechanisms affected by haploinsufficiency. In this approach, we took advantage of GeNets^30^, a computational framework for the analysis of protein-protein interactions, developed for the interpretation of genomic data. Our analysis allowed us to identify PPI clusters enriched for genes affected by CNVs and/or DNM in patients. Furthermore, GO enrichment analysis suggested distinct biological functions for four of these clusters.

Cluster 1 (**Figure 5a**) contains proteins involved in the Notch signaling pathway. Our data corroborate previous studies that confirm the central role of Notch pathway in the pathophysiology of CHD^51^ and highlights the shared contribution of CNVs and DNMs within the cluster. Cluster 3 (**Figure 5b**) contains proteins driving essentials processes in the development of the heart such as atrial septum and cardiac right ventricle morphogenesis as well as proteins playing significant role in the positive regulation of gene expression. These mechanisms has been well studied elsewhere^52^. Interestingly, three out of the seven candidate novel CHD genes (*WAC, ARID1B* and *KDMB5*) were found to be contributing to these two clusters. Cluster 8 (**Figure 5c**) showed enrichment for processes related with chromosome organization and DNA repair. Association between DNA repair and CHD is not well established thus far. Cluster 9 (**Figure 5d**) was found to be associated with microtubule organizing function. This biological process has been not described in the context of CHD, although an earlier report^53^ describes complex CHD among the phenotypes in individuals with 15q11.2 deletion syndrome, which involves the tubulin gamma complex protein 5.

Given the heterogenous data sources and the complex inheritance patterns often observed in patients with CHD, our study has limitations. Firstly, the patient data was collected from almost 200 different sources, and in many cases it was only possible to obtain data from CNV calls which had been already suggested to be pathogenic. Thus, we are aware that our patient data are incomplete because genome-wide CNV data are missing from a large part of the patient cohort. This is not the case for controls, for which genome-wide data was used. As a direct consequence, even though the difference between the rates of CNV deletions in controls and cases decreased dramatically after applying a quality control filtering step, a slight difference remained between both cohorts. In addition, the distribution of CNVs that overlap known microdeletion syndromes such as DiGeorge syndrome and Williams syndrome is overrepresented in the dataset. Similarly, the degree of phenotyping varied across the different studies, and often only basic phenotypic terms relating to CHD were available. This made it impossible to refine the diagnosis to a precise phenotypic class of CHD in many individuals. Nevertheless, the integrative approach allowed us to look for CHD associations in a binary fashion and will facilitate future studies and improve genotype-phenotype correlation for CHD subgroups.

In summary, we have performed an integrative analysis of CNVs, DNMs, o/e LOF ratio and expression during heart development amongst more than 10,000 CHD patients. Our analyses identify seven potential disease genes and mechanisms with novel association with CHD and strengthen previously reported associations.

## Materials and Methods

### Cohort description

Our cohort contains 7,958 CHD cases and 14,082 controls (see summary at **Supplemental Table 1**). Data from both affected and unaffected individuals were collected from 190 different CNV studies (**Supplemental Table 2**). Most of the CNV data included in the present study were assembled from public repositories, data available from literature as well as unpublished clinical data (see **Supplemental Information (Tables 1-3)** for a more detailed description). CHD phenotype information and genotyping platforms have been described in earlier publications (**Supplemental Table 2**). In addition, we built a dataset from the two largest DNM studies in CHD published thus far, which include a total of 2,489 parent-offspring trios.

### DNM analysis

The assembled DNM dataset was re-annotated using the Variant Effect Predictor (VEP version 90) tool. Based on the VEP annotation, we classified every variation into three major functional groups as follows: a) LOF variant (stop_gained, splice_acceptor, splice_donor, frameshift, initiator_codon, start_lost, conserved_exon_terminus), b) missense variant (stop_lost, missense, inframe_deletion, inframe_insertion, coding_sequence, protein_altering) and c) silent variant (synonymous). Variants with MAF > 0.01 in gnomAD database were excluded from the analysis. The rates of rare DNMs (MAF < 0.01) in both DNM studies^6,7^ were compared (Poisson test) for different variant consequence groups (PTV, missense and synonymous). No significant differences were found between the DNM rates for any of the evaluated groups (**Supplemental Table 9**). *De novo* mutation recurrence significance testing was performed to evaluate the impact of DNMs at gene level using the Mupit tool (https://github.com/jeremymcrae/mupit). By default, Mupit uses the sequence-specific mutation rate published by Samocha *et al*^54^. A second test, DeNovoWEST^16^, was used to assess gene-wise *de novo* mutation enrichment. DeNovoWEST assigns a mutation severity score (based on the variant consequences and the CADD score) to all classes of variants as a proxy of its deleteriousness. For each tested gene, the minimal *p-value* obtained from Mupit and DeNovoWEST was reported (*P*_*dnm*_). The corrected *P* value was computed using the Bonferroni method with n=18,272.

### CNV analysis

Only autosomal CNVs were included in the analysis. Also, smaller and longer CNVs were filtered out by applying a size cut-off of 5 Kb and 20 Mb as lower and upper limit, respectively. It has been demonstrated before that smaller and larger CNV calls tend to have a high rate of false positives^55,56^. We removed CNVs overlapping more than 50% of telomeres, centromeres and segmental duplication regions. In addition, we computed the internal CNV frequencies by counting the number of relative overlaps (>50% reciprocal overlap) on the CNVs control subset divided by the total number of controls. The internal MAF was computed for deletions and duplications subsets separately. Only CNVs with a minor allele frequency (MAF) < 0.01 in controls and overlapping ten or more CNV platform calling probes (Affymetrix Array 6.0 and Illumina Human660W-Quad) were considered for downstream analysis. Our analysis was focused only on CNV deletions. The distributions of the number of CNV deletions per individual within the case and control groups were compared (two-sided Wilcoxon rank sum test) to evaluate the impact of the quality control filtering step (**Supplemental Figure 5**). After filtering, 6,746 cases (3, 929 harbouring CNV deletions) and 14,024 controls (12,585 harbouring CNV deletions) remained for further analysis. A region-based permutation test (using PLINK version 1.07, test *‘--cnv-test-region --mperm 10000’*) was used on the filtered set to perform a case-control association analysis. For the gene-based permutation analysis, we reported both the ‘point-wise’ empirical p-value (EMP1) and the empirical adjusted p-value (EMP2), which controls the family-wise error rate (FWER) (http://zzz.bwh.harvard.edu/plink/perm.shtml). In addition to the gene-centered permutation testing, a similar region-based permutation analysis was performed to access enrichment in known CNV deletion syndromes. The region genomic coordinates and syndrome descriptions were downloaded from the *Database of genomic variation and phenotype in humans using Ensembl resources* (Decipher, https://decipher.sanger.ac.uk/disorders#syndromes/overview).

### CNV burden test on gene sets

A logistic regression-based burden test (‘cnv-enrichment-test’ in PLINK v1.7)^14^ was performed on different gene sets (known CHD genes (non-syndromic/syndromic/biallelic/monoallelic), developmental disorder genes, low observed/expected LOF ratio genes, **Supplemental Table 12**) using the rare CNV deletions passing the quality control and filtering stage. For every gene set examined, the binary phenotype (CHD case or control) was regressed on the number of genes disrupted by one or more CNVs. The averaged CNV size and the number of segments per individual were used as covariates into the model to control for potential differences between cases and controls as suggested by Raychaudhuri *et al*^14^. In addition, the PLINK implementation of this test was slightly modified by including a third (categorical) covariate, the sample study ID, since we have assembled the CNV data from different sources.

### Inferring differentially and highly expressed genes

Differentially expressed genes (DEGs) were identified by comparing the gene expression profile in heart to kidney and liver at matched time points. We used maSigPro R-package^57^ for inferring genes with dynamic temporal profiles from time-course transcriptomic data as previously described by Cardoso-Moreira *et al*^25^. As the input for maSigPro, we used the count per million matrix (CPM, output from EdgeR package) hosted in ArrayExpress (E-MTAB-6814). Genes which did not reach a CPM > 0.5 in at least five samples were excluded from the analysis. We ran maSigPro on the time scale measured in days post-conception using defaults parameters and only included time points with at least two biological replicates. A gene was selected as DEG if the *R*^*2*^ (goodness-of-fit) parameter was higher than 0.50 and Bonferroni corrected *P* < 0.01. The *R*^*2*^ parameter distinguish genes with clear expression trends from genes with ‘flat’ expression profile. **Supplemental Table 11** lists the final DEGs identified in the heart (*R*^*2*^ > 0.50). To assess the gene expression levels in the heart, the RPKM matrix was used. Gene expression levels were averaged among samples in the different development stages of the heart as follow: early development (4wpc-8wpc), maturation (9wpc-20wpc), infant/adult (newborn-adulthood). Genes were ranked based on the computed mean expression.

### Identification of CNV/DNM enriched PPI sub-clusters

A protein-protein interaction network was constructed using the GeNets framework and the information from InWeb and ConsensusPathDB protein-protein interaction databases. Nodes in the network correspond to proteins whereas edges represent their physical interactions. The network was strictly seeded with 492 candidate genes, those with a significant adjusted *metaP* < 0.05 (Benjamini-Hochberg’s false discovery rate, FDR). The PPI network was partitioned into overlapping sub-clusters using the in-built clustering method described in GeNets^30^. Only statistically significant sub-clusters (p-value < 0.05, permutation test) with at least 5 proteins were considered for further analysis. Finally, Gene Ontology enrichment analysis (Biological Process database 2018) of each identified sub-cluster was performed using the enrichr tool (http://amp.pharm.mssm.edu/Enrichr/).

## Data availability

The data used in this study have been already published. The reference for each individual study is shown in **Supplemental Table 1-3**. Upon request, we can provide the BED files.

## Funding resources

Funding for this study was provided by the DZHK and Kinderherzen consortia. BK is supported by a British Heart Foundation Personal Chair (CH/13/2/30154).

## Acknowledgements

We express our gratitude to the patients and their families for their participation in the analysed studies. We would like to thank the Genetic Association Information Network (GAIN) and dbGAP for making the data available. We would like to thank the Wellcome Trust Case Control Consortium (WTCCC) for making the data accessible https://www.wtccc.org.uk/info/access_to_data_samples.html. The Deciphering Developmental Disorders study presents independent research commissioned by the Health Innovation Challenge Fund (grant HICF-1009-003), a parallel funding partnership between the Wellcome Trust and the UK Department of Health, and the Wellcome Trust Sanger Institute (grant WT098051). The views expressed in this publication are those of the author(s) and not necessarily those of the Wellcome Trust or the UK Department of Health. We would like to thank the Pediatric Cardiac Genomics Consortium (PCGC) and dbGAP for making the data publicly available. This study was supported by the German Center for Cardiovascular Research (DZHK) partner sites Berlin, Kiel and Competence Network for Congenital Heart Defects, National Register for Congenital Heart Defects. This study makes use of data generated by the DECIPHER community. A full list of centres who contributed to the generation of the data is available from http://decipher.sanger.ac.uk and via email from. Funding for the project was provided by the Wellcome Trust. We would like to thank the Genome Aggregation Database (gnomAD) and the groups that provided exome and genome variant data to this resource. We thanks to Margarida C. Moreira and the Kaessmann Lab by making accessible the human RNA-Seq data and the support for the data analysis. We thanks Rasmus Wernersson and Federico de Masi for facilitating the use of the protein-protein interaction database, InWeb, in this work. We thanks the Lage Lab and Taibo Li for their support with GeNets. And finally, we would like to thank all data submitters and collaborators for their contributions.

