## Supplemental Figures for "Integrative analysis of genomic variants reveals new associations of candidate haploinsufficient genes with congenital heart disease"

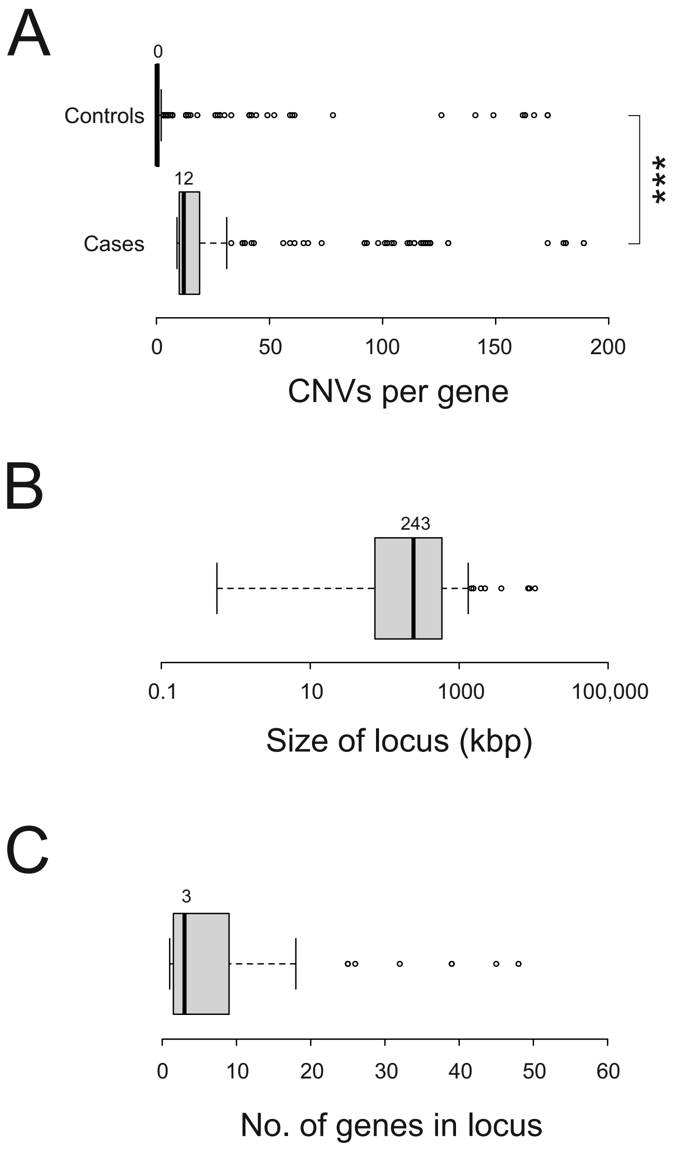


**Supplemental Figure 1. Distribution of overlapping CNVs, size and genes in 63 CHD loci. A) CNVs per gene.** Overlapping CNVs for each of the 528 significant candidate genes are shown as box-and-whiskers plots. Statistically significant difference was observed between the two distributions (Mann-Whitney test, ***: $P<0.001$). B) Size of loci in kilobase-pairs (kbp). C) Number of genes per locus. Median values are shown above each box.

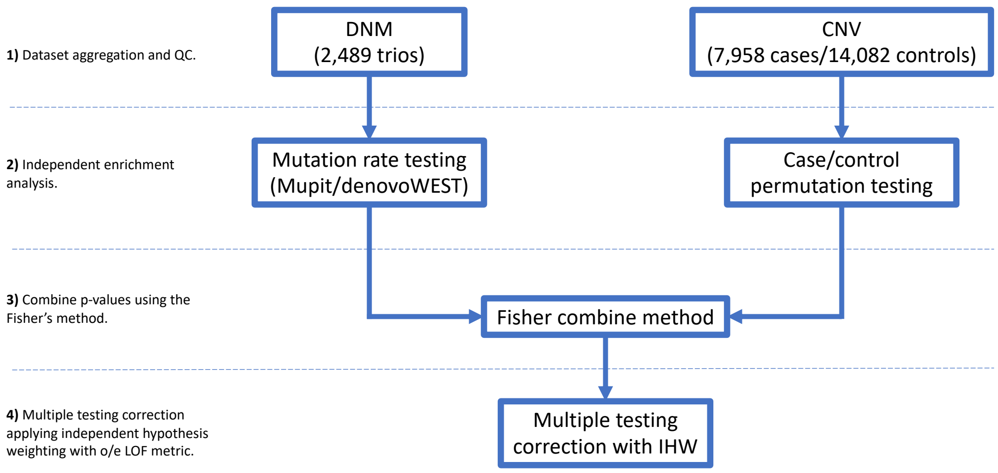


**Supplemental Figure 2. Statistical framework to discover novel candidate CHD genes by integrating DNM and CNV deletions.** The workflow follows four major steps: 1) Data aggregation and quality control of both DNM data and CNV data (shown the number of cases/controls passing the quality control filters), 2) DNM rate-based enrichment testing and CNV deletions case/control association analysis at gene level are performed independently, 3) the results are combined using the Fisher method and 4) P-values are Bonferroni corrected using the Independent Hypothesis Weighting method (IHW). As independent covariate for the IHW method, the o/e LOF ratio upper bound fraction (LOEUF) was used.

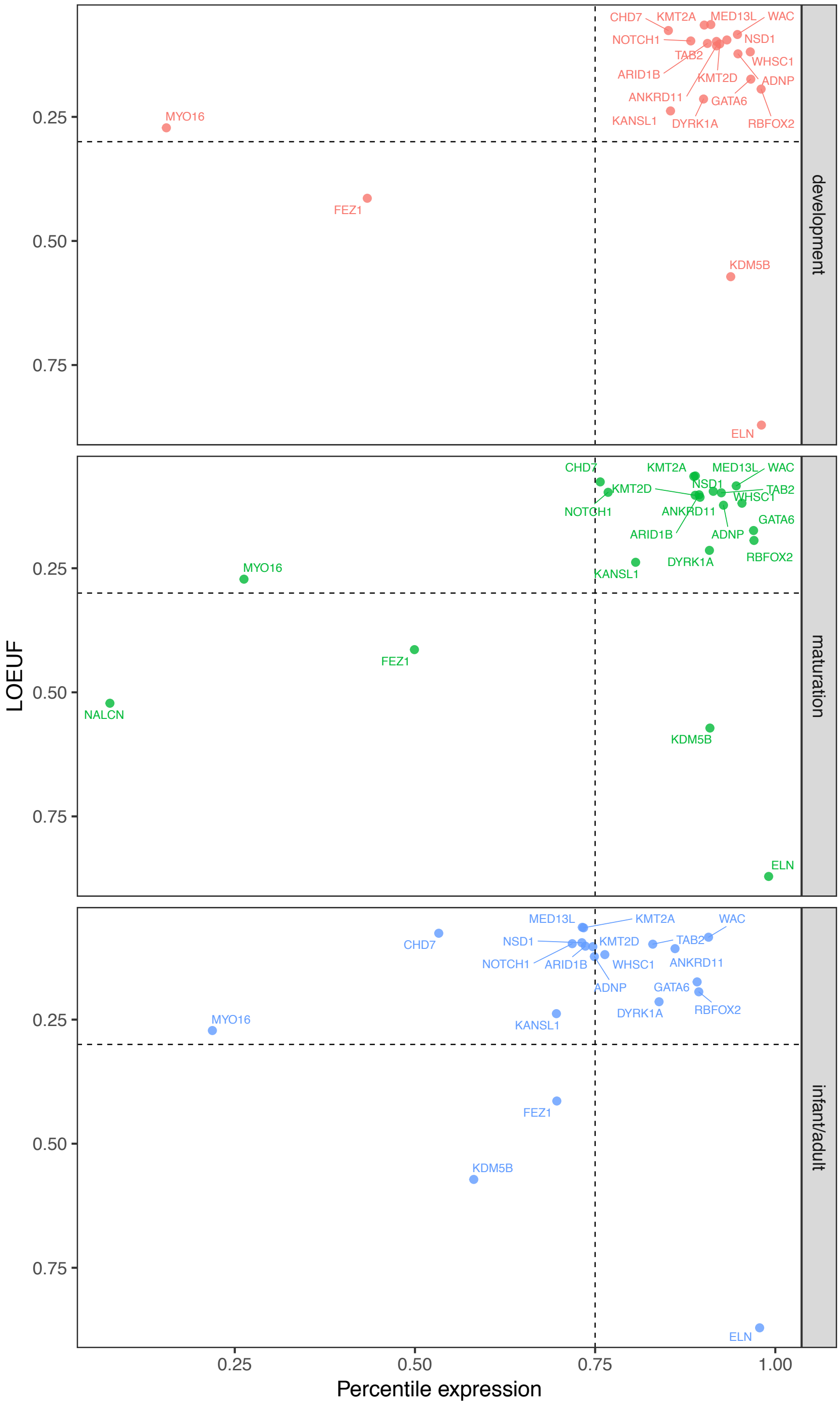


**Supplemental Figure 3. Heart expression pattern of the 21 significant genes at different heart development stages.** Panels show three different heart development stages: early development (red), maturation (green) and infant/adult (blue). The x-axis denotes the percentile rank of heart expression in the heart. The y-axis denotes the o/e LOF ratio upper bound fraction (LOEUF) from gnomAD. Dashed lines denote the threshold for highly expressed genes (expression rank >= 0.75) and highly LOF constrained genes (LOEUF <= 0.30).

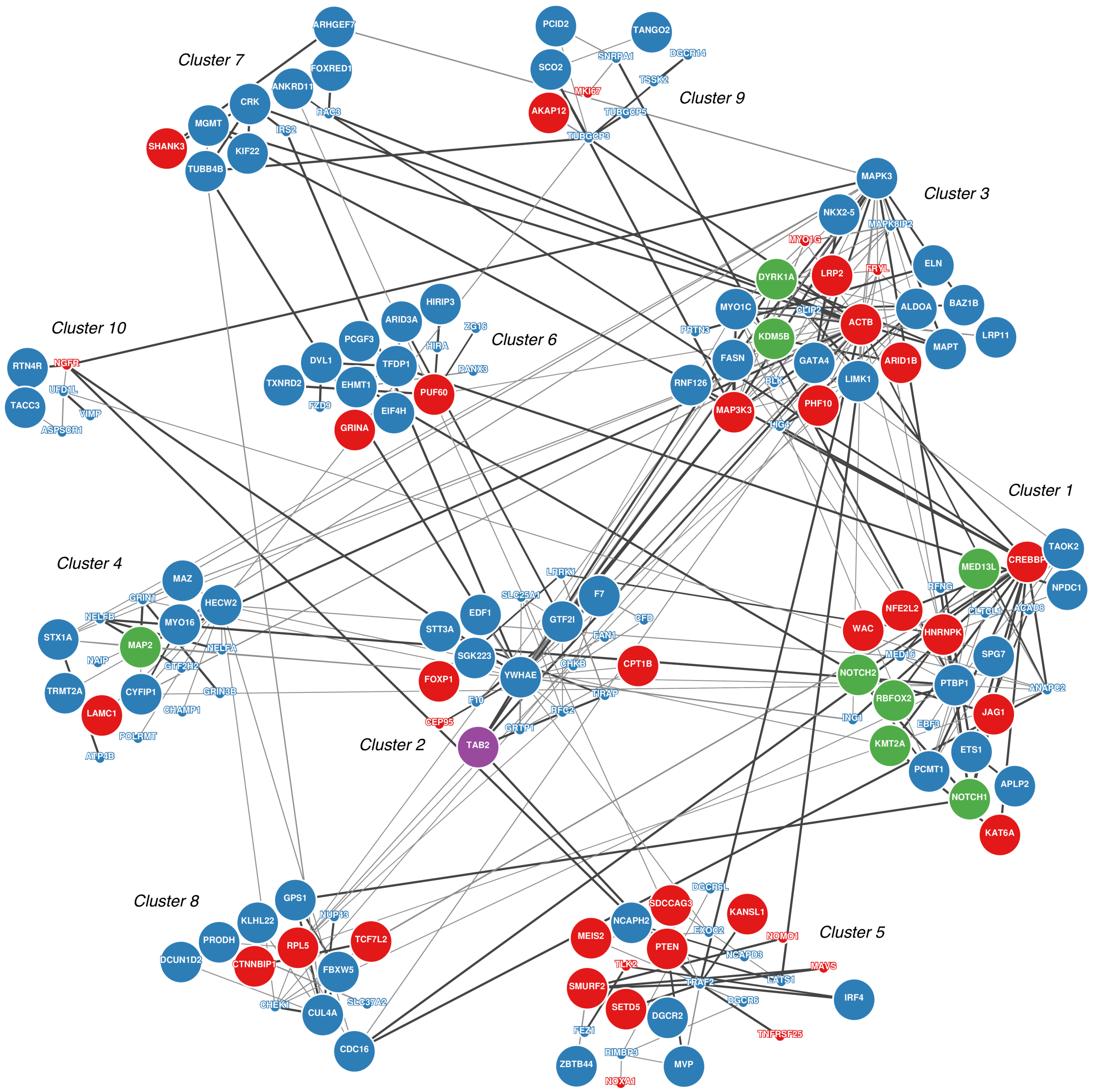


**Supplemental Figure 4. The functional network enriched for proteins encoded by genes affected by CNVs and/or DNMs associated with CHD.** Ten sub-clusters were identified using GeNets. Proteins are shown as nodes, interactions as edges. Enrichment for CNVs (blue), DNMs (green) or both independently (purple) are highlighted. Proteins with no specific enrichment for CNV and/or DNMs but with B-H adjusted *metaP* < 0.05 are highlighted in red. The size of the circles denotes if the gene was found significantly highly and/or differentially expressed in the heart (large circles: significant expression; small circles: non-significant).

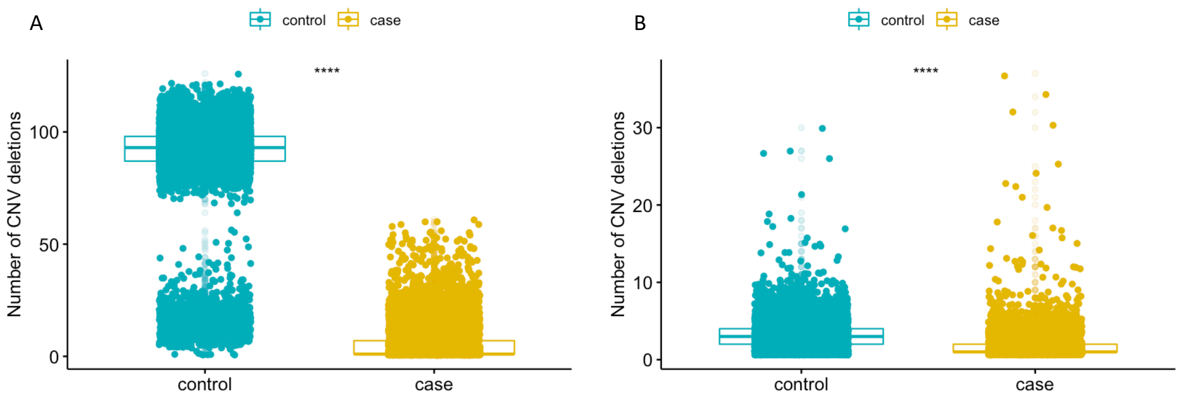


**Supplemental Figure 5.** Distribution of the number of CNV deletions per individual in both control and CHD case cohorts before (A) and after (B) applying the quality control filtering approach. Differences between the distributions were tested using a two-sided Wilcoxon rank sum test. ****: P<0.0001.
